# A transient time window for early predispositions in newborn chicks

**DOI:** 10.1101/623439

**Authors:** Elisabetta Versace, Morgana Ragusa, Giorgio Vallortigara

**Affiliations:** Department of Biological and Experimental Psychology, School of Biological and Chemical Sciences, Queen Mary University of London, London (UK); Center for Mind/Brain Sciences, University of Trento, Rovereto (Italy)

**Keywords:** Early predispositions, sensitive period, changes in speed, early development, newborn chicks, *Gallus gallus*, neonates

## Abstract

Neonates of different species are born with a set of predispositions that influence their early orienting responses toward the first stimuli encountered in their life. Human neonates and domestic chicks exhibit several similarities in the predisposition for attending to objects that move with speed changes, face-like stimuli and biological motion. Although early predispositions are connected to physiological development, little is known on the temporal course of early predispositions (whether they are stable or change in time) and on their genetic basis. To address these issues, we tested the preference for objects that change in speed vs. linear motion in three chicken breeds (Padovana, Polverara and Robusta maculata) within one day after hatching and three days after hatching. We found that the predisposition to preferentially attend to changes in speed is fixed at the species level on the first day of life and that it disappears by day three. These results indicate the existence of a short and transient time window of early predispositions that does not depend on visual experience.

## Introduction

For human adults, objects that move with visible speed changes appear animate, as if they move through an internal energy source^1^. Experiments with very young and inexperienced subjects show that the relevance of dynamic changes in speed does not need to be learned. In fact, both human neonates^2^ and visually naïve chicks^1^ are more attracted to objects that move changing speed compared to objects that move linearly. These data suggest that this early unlearned preference might be widespread among vertebrate species. The aim of this work is to shed light on the temporal course and genetic basis of this predisposition.

The preference for speed changes has been described as a predisposition to attend to “animacy” cues, the properties associated with living beings^3,4^. Other animacy cues that are attractive for neonates include face-like stimuli^4–9^, biological motion^10,11^ and self-propulsion^12^. The initial orienting response towards animate objects, in turn, enhances learning and social interactions. This phenomenon has been described in domestic chicks, that quickly develop affiliative responses towards the first objects they are exposed to (especially for moving objects), through the mechanism of filial imprinting^13–16^. Imprinting can also modulate predisposed responses^17^. While chicks are a convenient model to investigate early predispositions – because they are a precocial^18^ species that can be easily tested in the absence of previous visual experience – human babies show striking similarities in the response to the stimuli preferentially attended and approached by chicks (^6,10–12,19–25^). Interestingly, both human neonates at high familial risk of autism^23^ and chicks exposed to drugs that increase the risk of developing autism in humans^24,25^ exhibit an impairment (or a temporal shift) in early predispositions for animacy. This is another piece of evidence that points at a possible conserved origin of early predispositions in vertebrates. Comparative studies are important to clarify the origins and neurobiological mechanisms of early predispositions.

Some works have started to shed light on the temporal course of early predispositions. Previous studies on chicks’ predisposition to preferentially approach a stuffed-hen vs. a scrambled version of the same stimulus showed that 24 hours after motor stimulation (walking on a wheel) chicks exhibited a much stronger preference for the stuffed-hen than after 2 hours from motor stimulation^26,27^. The preference for the stuffed hen is linked to the head and neck region of the model, suggesting that this predisposition is connected to the preference for the inverted triangular shape of the face-like stimuli^5,26^. While even human foetuses have been shown to preferentially orient towards this pattern^28^, it is known that the preference for face-like patterns is not stable during development, declining at around two months of age^29–31^. Shultz and colleagues^32^ made the interesting point of a general pattern of transitions from reflexive behaviours to volitional actions in human infants. A working hypothesis is that early predispositions are transient not only in human neonates but in other species as well, and that they decline at an age in which volitional control in approach responses takes over. Such a mechanism would be fixed at the species level, showing little genetic variability.

The temporal course of the predisposition for changes in speed has not been previously investigated. To address this issue, we tested chicks in the first day after hatching (post-hatching day 0, p0) and three days after hatching (post-hatching day 2, p2). Previous studies showed that while the very first orienting responses to the stuffed hen are fixed at the species level, the subsequent responses of sustained attention have a genetic basis that differs between breeds^6^. To investigate whether the predisposition to orient towards changes in speed and its temporal course have genetic variability, we conducted the study in three breeds of domestic chickens that have been kept genetically isolated for 20 years in the same farm: Padovana, Polverara and Robusta maculata. This arrangement ensures that differences between breeds are not due to environmental factors.

## Methods

### Ethics statement

All experiments comply with the current Italian and European Union laws for the ethical treatment of animals and the experimental procedures were approved by the Ethical Committee of University of Trento and licensed by the Italian Health Ministry (permit number 1138/2015 PR).

### Subjects and rearing conditions

Fertilized eggs of three breeds of domestic fowl (*Gallus gallus*) kept genetically isolated for 20 years in the conservation programme CO.VA^33,34^ were obtained from the Agricultural High School “Duca degli Abruzzi”, Padova (Italy). Further details on this population, breeds and their early predisposition are available in previous publications ^6,35^. Eggs were incubated in darkness at a constant temperature of 37.7 °C, humidity 40% until 3 days before hatching (day 17 of incubation). Three days before hatching, the humidity was increased at 60% and eggs of each breed were located on the same tray for hatching. Until the moment of the test, chicks were kept in constant darkness inside the incubator in a dark room, making sure they did not receive any visual stimulation until the moment of the test. Each chick was tested only once.

In Experiment 1, we tested 117 domestic chicks (*Gallus gallus*) and analysed the performance of the 88 chicks that moved during the test (it was possible to calculate a preference index only for these chicks): 31 chicks of Padovana breed (PD), 30 chicks of Polverara breed (PL), 27 chicks of Robusta maculata breed (RB). Chicks were tested at day post-hatching 0 (p0, 12-24 hours after hatching) and were kept in the incubator until the moment of the test.

In Experiment 2, we tested 77 chicks and analysed the performance of the 51 domestic chicks that moved during the test (it was possible to calculate a preference index only for these chicks): 31 chicks of Padovana breed (PD) and 20 chicks of Polverara Breed (PL). Chicks were tested at day post-hatching 2 (p2, 3 days after hatching) and were kept in the incubator until the moment of the test. In this experiment, only two breeds were used because there no more eggs of the Robusta breed were available.

After the test, chicks were housed in groups and maintained with food and water available ad libitum in standard rearing conditions until donated to local farmers.

### Apparatus

The test apparatus was a rectangular black box (85 cm length × 62 cm width × 58 cm height). At the opposite sides of the apparatus, two monitors (LCD Monitor BenQ XL2410T, resolution 1920×1080 pixels, refresh frequency 120 Hz, 48.7 cm width × 36.5 height) played the stimuli. One monitor showed the stimulus with changes in speed (animate stimulus), the other one showed the linearly moving stimulus (see section Stimuli), se Figure 1.

**Figure 1.**
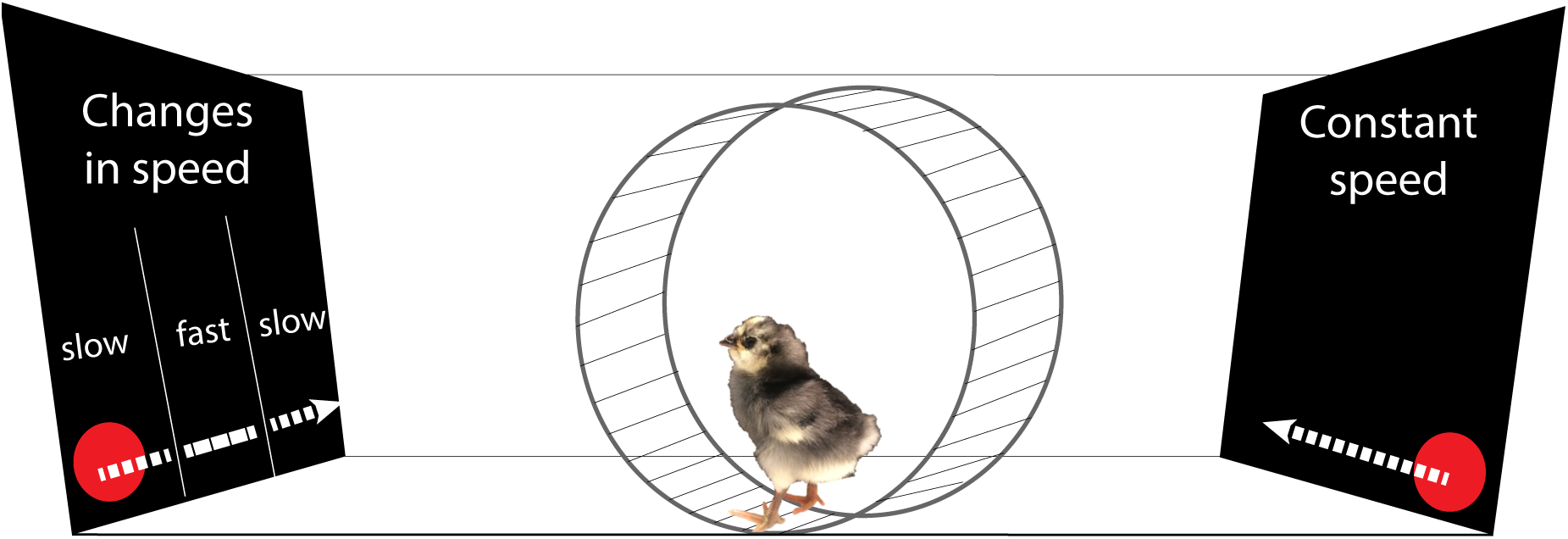
Schematic representation of the experimental setting. We measured chicks’ approach responses (distance run in a wheel) towards a stimulus that moved changing in speed (the “animate” stimulus) and an object moving linearly with constant speed (the “non-animate” stimulus).

A running wheel (30 cm diameter, 13 cm width, covered with 1 cm of opaque foam on both sides) was located in the middle of the apparatus at a distance of 40 cm from each monitor). Above the wheel, a video-camera (Windows Lifecam) at a distance of 77 cm from the bottom of the running wheel recorded the experimental sessions. We recorded the centimetres run by the chicks in the wheel in both directions using magnetic sensors located on the sides of the wheel. In the running wheel, subjects could easily change the direction of movement. The experiment was run in darkness except for the stimuli played on the monitors.

### Stimuli

We played the same stimuli used in a previous study for the measurement of spontaneous preferences for speed changes in visually naïve chicks^1^. Briefly, the stimuli consisted of video animations of a red disk (3 cm diameter) on a black background moving horizontally. A horizontal grey band was used as a floor on which the stimulus moved. The 30 cm wide screen was bounded by lateral grey bars at the sides (3 cm wide), but placed on the screen in order to occupy only 2.5 cm of visible screen space, so that the visible trajectory of the red stimulus was 24.6 cm long, see ^1^ for further details).

The two stimuli were presented simultaneously on the opposite monitors. The stimulus always entered the visible scene already in motion and then disappeared from view while still moving. One stimulus always exhibited visible speed changes along its trajectory (the “animate stimulus”), whereas the other stimulus moved at a constant speed (the “not animate stimulus”). The not animate stimulus moved at constant 4.64 cm/s speed. The animate stimulus had an initial speed of 3.37 cm/s, at one-third of its way it abruptly accelerated to 19.64 cm/s up to two-thirds of the route, where abruptly decelerated at the initial speed. Both stimuli moved at the same average speed. Previous control experiments showed that the response to change in speed, and not a generic response to change, drive the preference exhibited by chicks^1^. The stimuli appeared and disappeared simultaneously, and appeared again after one second of delay on the same side from where they disappeared. The stimuli were presented to the chick as moving, and continued looping for the whole duration of the test (30 minutes).

### Test Procedure

At the beginning of the test, each chick was taken from the incubator and transported to the test room, constantly kept in dark. Each individual was placed in the running wheel facing the long side of the apparatus, oriented with the right eye facing the right screen. In this way, the chick could simultaneously see both stimuli.

In the wheel, the chick could walk towards each stimulus, and the distance run (in centimetres) by every chick towards each side (right/left) was recorded every 10 minutes (at 600, 1200, 1800 seconds), for 3 time periods (30 minutes overall). Besides investigating the overall motor activity as number of centimetres run, we calculated a preference index for the animate stimulus as

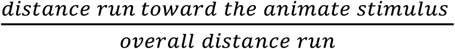

In this index, 1 indicates a complete preference for the animate stimulus, 0 a complete preference for the not animate stimulus and 0.5 no preference.

### Statistical Analysis

To analyse the spontaneous preference for the animate stimulus, we performed an ANOVA using Breed (Padovana, Polverara, Robusta) as between subjects variable and Time (minutes 0-10, 10-20, 20-30) as a within subjects variable. For the comparison between chicks tested at day post-hatching 0 (Experiment 1) and day post-hatching 2 (Experiment 2), we performed an ANOVA using Breed and Day post-hatching (p0, p2) as between subjects variables and Time (minutes 0-10, 10-20, 20-30) as within subjects variable. We used one sample t-tests to assess departures from the chance level (0.5).

Because the assumptions of parametrical tests were not satisfied, to analyse the differences between Breeds in motor activity (centimetres run) we used the Kruskall-Wallis test with Breed as independent variable. We used the Mann-Whitney test for post-hoc comparisons.

For all tests, significance was set to p<0.05. Statistical analyses were performed with the software R (version 3.5.2).

## Results

### Spontaneous preference for the animate stimulus

In Experiment 1 (test at post-hatching day 0), we observed no significant main effect of Breed (F_2,85_=0.265, p=0.768) and Time (F_2,170_=0.279, p=0.757) but a significant interaction Breed x Time (F_4,170_=4.227, p=0.003), see Figure 2a. Since all data points were located above the chance level and there was no main effect of Breed, we also tested the overall performance of the population against the chance level. The overall population exhibited a significant preference for the animate stimulus (t_87_=2.917, p=0.004, Mean=0.590, SD=0.289).

**Figure 2.**
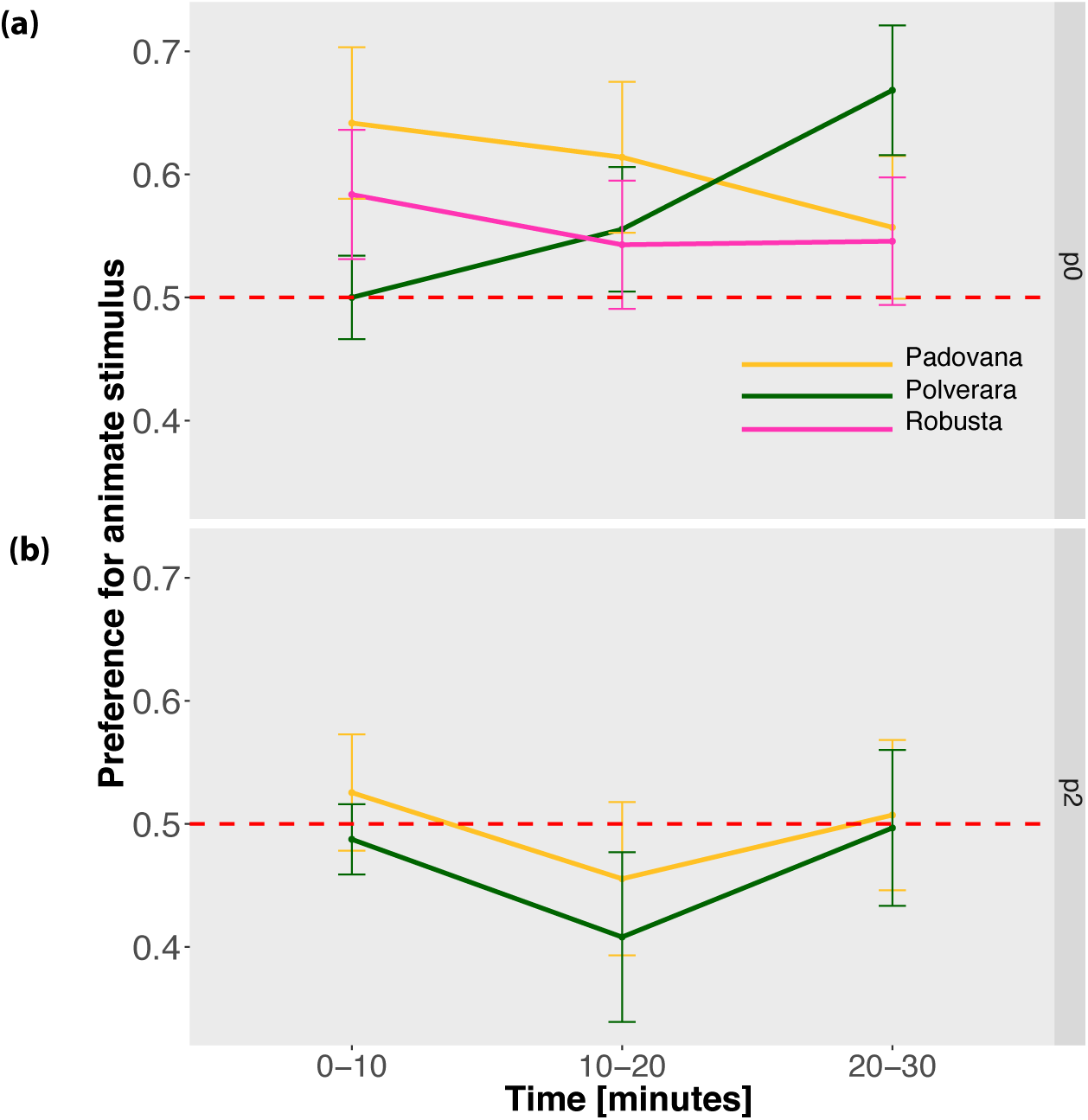
The lines show for each Breed the Mean +/− Standard error of the mean, for the preference for the animate stimulus in (a) Experiment 1 and (b) Experiment 2 across the three time periods: minutes 0-10, 10-20 and 20-30.

In Experiment 2 (test at post-hatching day 2), we observed no significant main effect of Breed (F_1,49_=0.255, p=0.616) and Time (F_2,98_=01.458, p=0.238) and a non significant interaction Breed × Time (F_2,98_=5.639, p=0.057), see Figure 2b. The overall population exhibited no significant difference from the chance level (t_50_=-0315, p=0.754, Mean=0.489, SD= 0.313).

### Motor activity

In Experiment 1 we observed a significant difference in motor activity between breeds (Kruskal-Wallis chi-squared=16.071, p<0.001; see Figure 3a. Padovana chicks ran significantly more than Polverara (U=667.5, p=0.004) and Robusta (U=174, p<0.001) chicks, while there was no significant difference between Robusta and Polverara (U=350.5, p=0.388). In Experiment 2 there was no significant difference between Padovana and Polverara chicks (Kruskal-Wallis chi-squared=2.265, p=0.132), although Padovana had a trend for running more than Polverara (see Figure 3b), as in Experiment 1. Overall, chicks ran more in Experiment 2 than in Experiment 1 (U=2892, p=0.005).

**Figure 3.**
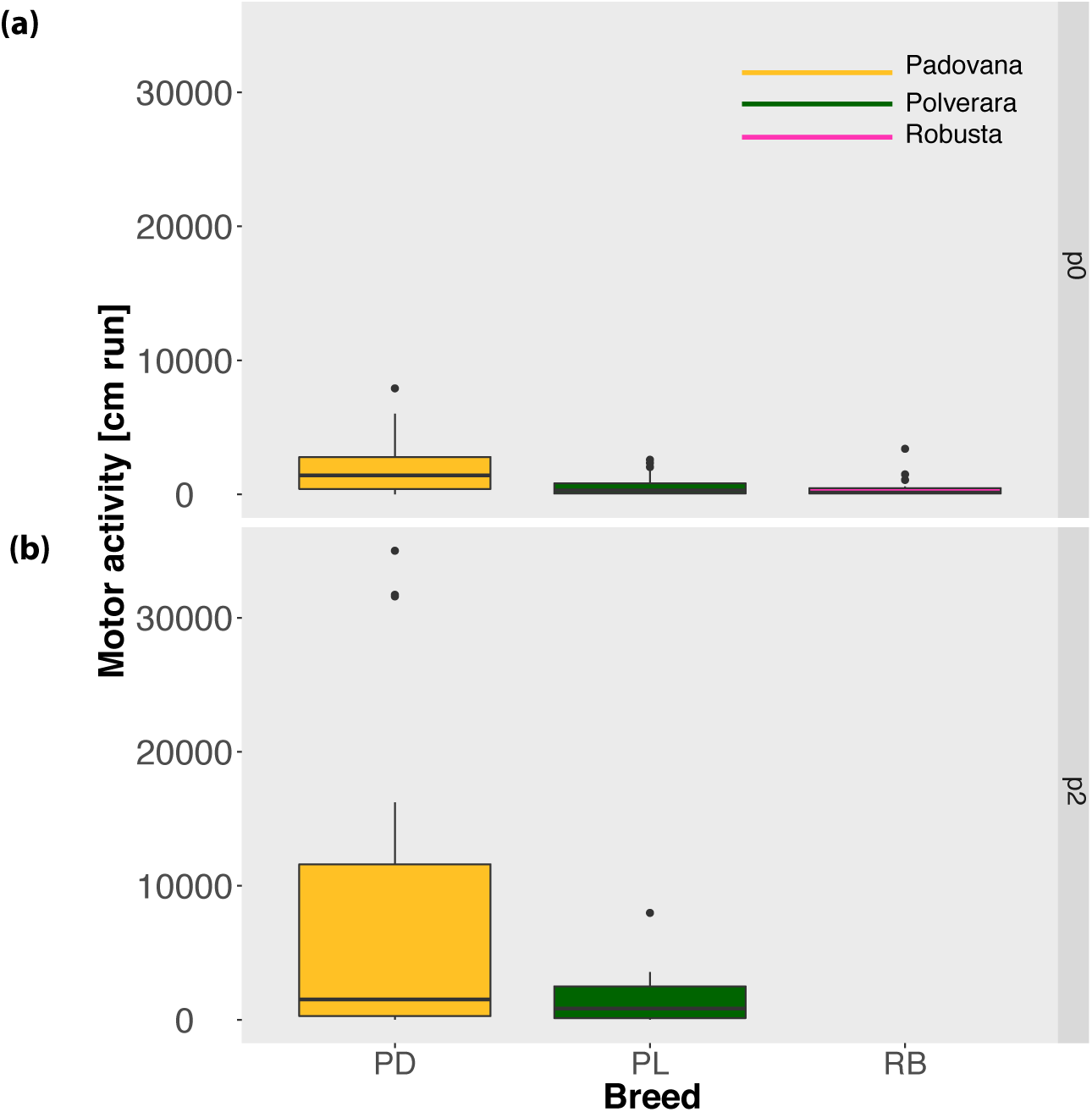
The boxplots show for each Breed the median, interquartile range, minimum, maximum and outliers for the motor activity (centimetres run) in (a) Experiment 1 and (b) Experiment 2.

## Discussion

Studying the early responses of chicks of three different chicken breeds (Padovana, Polverara, Robusta maculata), we found that the predisposition to preferentially approach an object that moves changing speed vs. a linearly moving object is a trait shared at the species level in the first day of life (post-hatching day 0). This preference, though, is not present two days later (p2). This decline is in line with other reports of human neonates that lose an initial predisposition^8,9,21,22^ (in the absence of previous experience) for orienting towards face-like stimuli while moving to a voluntary control of the attention (reviewed here^32^), as well as a peak in the predisposition to approaching a stuffed hen vs. control stimuli in chicks at around 24 hours of age^26,27^.

While the study of predispositions in human neonates is constrained by practical and ethical issues, it is possible to deprive domestic chicks from any visual experience until the moment of the test to understand whether the course of early visual predisposition depends on experience. Here, we kept chicks visually naïve until the moment of the test to excluded that the decline in the predisposition is due to experience. The increase of general motor activity (distance run) with time, when the predisposition was not apparent anymore, constitutes strong evidence that the decline in the predisposition is not due to a reduced interest in approaching the stimuli at post-hatching day 2. On the contrary, older chicks were more motivated in approaching the experimental stimuli, in spite of the lack of a predisposed preference for the object that changed speed. These findings suggest that the decline of the predisposition to attend to speed changes does not depend on experience but rather on developmental processes of maturation. It has been previously shown that, although chicks are a precocial species with developed motor and sensory abilities at birth, their nervous system undergoes a maturation process in the first stages of life^36^, similarly to what happens in human neonates^32^.

The implications of delayed or impaired early predispositions for the physiological development have still to be clarified. Yet, a generalised pattern of absent or delayed predispositions has been already documented both in human neonates at high risk of autism^23^ and newborn chicks exposed to autism-inducing drugs^24,25^. The domestic chick appears to be an ideal model for studying early predispositions for several reasons: the study of early predispositions is easy in avian species, whose experience can be manipulated more easily from embryonic development; extensive parallels in early predispositions between neonate chicks and humans have been extensively documented, and our results confirm that early predispositions are widespread among vertebrate species and fixed in the domestic chicken; both physiological and pathological patterns appear to converge in human neonates and chicks in the first hours after birth; our findings point towards a basis of early predispositions that might not depend on visual experience, and whose neurobiological basis has already clarified several target areas in the chick brain such as the lateral septum, thee preoptic area and the nucleus taeniae of the amygdala^37–41^. Further studies should clarify the developmental mechanisms (e.g. the role of the thyroid hormones T_3_ and Dio2^42,43^) that sustain the transient presence of early predispositions in neonate animals.

## Author Contributions

E.V., and G.V. conceived the project; E.V. designed the experiments; E.V. and M.R. conducted the experiments, E.V. and M.R. analysed the data; E.V. drafted the manuscript; all the authors contributed to the manuscript and gave final approval for publication.

## References

1. Rosa-Salva, O., Grassi, M., Lorenzi, E., Regolin, L. & Vallortigara, G. Spontaneous preference for visual cues of animacy in naïve domestic chicks: the case of speed changes. Cognition 157, 49–60 (2016).

2. Di Giorgio, E., Lunghi, M., Simion, F. & Vallortigara, G. Visual cues of motion that trigger animacy perception at birth: The case of self-propulsion. Dev. Sci. 1–12 (2016). doi:10.1111/desc.12394

3. Di Giorgio, E. et al. Filial responses as predisposed and learned preferences: Early attachment in chicks and babies. Behav. Brain Res. 325, 90–104 (2017).

4. Rosa Salva, O., Mayer, U. & Vallortigara, G. Roots of a social brain: Developmental models of emerging animacy-detection mechanisms. Neurosci. Biobehav. Rev. 50, 150–168 (2015).

5. Rosa-Salva, O., Regolin, L. & Vallortigara, G. Faces are special for newly hatched chicks: evidence for inborn domain-specific mechanisms underlying spontaneous preferences for face-like stimuli. Dev. Sci. 13, 565–77 (2010).

6. Versace, E., Fracasso, I., Baldan, G., Dalle Zotte, A. & Vallortigara, G. Newborn chicks show inherited variability in early social predispositions for hen-like stimuli. Sci. Rep. 7, 40296 (2017).

7. Johnson, M. H. & Horn, G. Development of filial preferences in dark-reared chicks. Anim. Behav. 36, 675–683 (1988).

8. Goren, C. C., Sarty, M. & Wu, P. Y. K. Visual Following and Pattern Discrimination of Face-like Stimuli by Newborn Infants. Pediatrics 56, 544–549 (1975).

9. Buiatti, M. et al. Cortical route for facelike pattern processing in human newborns. Proc. Natl. Acad. Sci. U. S. A. 116, 1–6 (2019).

10. Vallortigara, G., Regolin, L. & Marconato, F. Visually inexperienced chicks exhibit spontaneous preference for biological motion patterns. PLoS Biol. 3, e208 (2005).

11. Simion, F., Regolin, L. & Bulf, H. A predisposition for biological motion in the newborn baby. Proc. Natl. Acad. Sci. U. S. A. 105, 809–13 (2008).

12. Mascalzoni, E., Regolin, L. & Vallortigara, G. Innate sensitivity for self-propelled causal agency in newly hatched chicks. Proc. Natl. Acad. Sci. U. S. A. 107, 4483–5 (2010).

13. Miura, M. & Matsushima, T. Biological motion facilitates imprinting. Anim. Behav. 116, 171–180 (2016).

14. Versace, E., Martinho-Truswell, A., Kacelnik, A. & Vallortigara, G. Priors in Animal and Artificial Intelligence: Where Does Learning Begin? Trends Cogn. Sci. 22, 963–925 (2018).

15. Bolhuis, J. J. Early learning and the development of filial preferences in the chick. Behav. Brain Res. 98, 245–52 (1999).

16. Vallortigara, G. & Versace, E. Filial Imprinting. Encyclopedia of Animal Behavior 1943–1948 (2018).

17. Versace, E., Schill, J., Nencini, A. M. & Vallortigara, G. Naïve chicks prefer hollow objects. PLoS One 11, (2016).

18. Versace, E. in Encyclopedia of Animal Cognition and Behavior 1–3 (Springer International Publishing, 2018). doi:10.1007/978-3-319-47829-6_459-2

19. Mascalzoni, E., Regolin, L., Vallortigara, G. & Simion, F. The cradle of causal reasoning: Newborns’ preference for physical causality. Dev. Sci. 16, 327–335 (2013).

20. Bardi, L., Regolin, L. & Simion, F. Biological motion preference in humans at birth: Role of dynamic and configural properties. Dev. Sci. 14, 353–359 (2011).

21. Simion, F. & Di Giorgio, E. Face perception and processing in early infancy: Inborn predispositions and developmental changes. Front. Psychol. 6, 1–11 (2015).

22. Morton, J. & Johnson, M. H. CONSPEC and CONLERN: A two-process theory of infant face recognition. Psychol. Rev. 98, 164–181 (1991).

23. Di Giorgio, E. et al. Difference in Visual Social Predispositions Between Newborns at Low- and High-risk for Autism. Sci. Rep. 6, 26395 (2016).

24. Lorenzi, E., Pross, A., Rosa-Salva, O., Versace, E. & Sgadò, P. Embryonic Exposure to Valproic Acid impairs Social Predispositions for Dynamic Cues of Animate Motion in Newly-Hatched Chicks. Front. Physiol. (2019). doi:http://dx.doi.org/10.1101/412635.

25. Sgadò, P., Rosa-Salva, O., Versace, E. & Vallortigara, G. Embryonic Exposure to Valproic Acid Impairs Social Predispositions of Newly-Hatched Chicks. Sci. Rep. 8, 5919 (2018).

26. Johnson, M. H. & Horn, G. Development of filial preferences in dark-reared chicks. Anim. Behav. 36, 675–683 (1988).

27. Johnson, M. H., Bolhuis, J. J. & Horn, G. Interaction between acquired preferences and developing predispositions during imprinting. Anim. Behav. 33, 1000–1006 (1985).

28. Reid, V. M. et al. The Human Fetus Preferentially Engages with Face-like Visual Stimuli. Curr. Biol. 27, 1825–1828.e3 (2017).

29. Davison, A. et al. Formin Is Associated with Left-Right Asymmetry in the Pond Snail and the Frog. Curr. Biol. 26, 654–660 (2016).

30. Simion, F., Valenza, E., Umiltà, C. & Dalla Barba, B. Preferential orienting to faces in newborns: a temporal-nasal asymmetry. J. Exp. Psychol. Hum. Percept. Perform. 24, 1399–405 (1998).

31. Johnson, M. H., Dziurawiec, S., Ellis, H. & Morton, J. Newborns’ preferential tracking of face-like stimuli and its subsequent decline. Cognition 40, 1–19 (1991).

32. Shultz, S., Klin, A. & Jones, W. Neonatal Transitions in Social Behavior and Their Implications for Autism. Trends Cogn. Sci. 22, 452–469 (2018).

33. Zanetti, E., De Marchi, M., Abbadi, M. & Cassandro, M. Variation of genetic diversity over time in local Italian chicken breeds undergoing in situ conservation. Poult. Sci. 90, 2195–201 (2011).

34. Zanetti, E., De Marchi, M., Dalvit, C. & Cassandro, M. Genetic characterization of local Italian breeds of chickens undergoing in situ conservation. Poult. Sci. 89, 420–7 (2010).

35. Versace, E., Ragusa, M. & Pallante, V. Conserved abilities of individual recognition and genetically modulated social responses in young chicks (Gallus gallus).

36. Rogers, L. J. The Development of Brain and Behaviour in the Chicken. (CAD International, 1995).

37. Mayer, U., Rosa-Salva, O., Loveland, J. & Vallortigara, G. Selective response of the nucleus taeniae of the amygdala to a naturalistic social stimulus in visually naive domestic chicks. submitted

38. Lorenzi, E., Mayer, U., Rosa-Salva, O. & Vallortigara, G. Dynamic features of animate motion activate septal and preoptic areas in visually naïve chicks (Gallus gallus). Neuroscience 354, 54–68 (2017).

39. Mayer, U., Rosa-Salva, O., Morbioli, F. & Vallortigara, G. The motion of a living conspecific activates septal and preoptic areas in naive domestic chicks (*Gallus gallus*). Eur. J. Neurosci. 45, 423–432 (2017).

40. Mayer, U., Rosa-Salva, O. & Vallortigara, G. First exposure to an alive conspecific activates septal and amygdaloid nuclei in visually-naïve domestic chicks (*Gallus gallus*). Behav. Brain Res. 71–81 (2017).

41. Mayer, U., Rosa-Salva, O., Lorenzi, E. & Vallortigara, G. Social predisposition dependent neuronal activity in the intermediate medial mesopallium of domestic chicks (*Gallus gallus domesticus*). Behav. Brain Res. 310, 93–102 (2016).

42. Takemura, Y. et al. Gene expression of Dio2 (thyroid hormone converting enzyme) in telencephalon is linked with predisposed biological motion preference in domestic chicks. Behav. Brain Res. 2, 25–30 (2018).

43. Yamaguchi, S. et al. Thyroid hormone determines the start of the sensitive period of imprinting and primes later learning. Nat. Commun. 3, 1081 (2012).

